# Stronger connectivity of the resident gut microbiome lends resistance to invading bacteria

**DOI:** 10.1101/261750

**Authors:** Cristina M. Herren, Michael Baym

## Abstract

Bacterial infection in the gut is often due to successful invasion of the host microbiome by an introduced pathogen. Ecological theory indicates that resident community members and their interactions should be strong determinants of whether an invading taxon can persist in a community. In the context of the gut microbiome, this suggests colonization resistance against newly introduced bacteria should depend on the instantaneous bacterial community composition within the gut and interactions between these constituent members. Here we develop a mathematical model of how metabolite-dependent biotic interactions between resident bacteria mediate invasion, and find that stronger biotic connectivity from metabolite cross-feeding and competition increases colonization resistance. We then introduce a statistical method for identifying invasive taxa in the human gut, and show empirically that greater connectivity of the resident gut microbiome is related to increased resistance to invading bacteria. Finally, we examined patient outcomes after fecal microbiota transplant (FMT) for recurring *Clostridium difficile* infection. Patients with lower connectivity of the gut microbiome after treatment were more likely to relapse, experiencing a later infection. Thus, simulation models and data from human subjects support the hypothesis that stronger interactions between bacteria in the gut repel invaders. These results demonstrate how ecological invasion theory can be applied to the gut microbiome, which might inform targeted microbiome manipulations and interventions. More broadly, this study provides evidence that low connectivity in gut microbial communities is a hallmark of community instability and susceptibility to invasion.

## Introduction

Human infection is often due to the successful invasion of harmful bacteria in the host-associated microbiome (Nizet et al. 2001, Bel et al. 2017). Although the human body is frequently exposed to harmful invaders (for example, pathogens on food [Berger et al. 2010] or household surfaces [Flores et al. 2013]), few introductions result in disease. However, pathogens comprise only a small subset of all potentially invasive bacteria within the human gut microbiome (David et al. 2014). The resident microbiome plays a substantial role in mediating resistance to invaders, but the mechanisms of action are only partially understood (Baumler and Sperandio 2016).

Ecological invasions progress through distinct stages, beginning with transport and population expansion, and culminating in their impact on ecosystems (Sakai et al. 2001). These same stages occur when newly introduced bacteria colonize the human microbiome (Bosch et al. 2013, Kc et al. 2017). Most ecological invasions are unsuccessful (Williamson 2006). This is also observed in invasions in the human microbiome; although humans are constantly exposed to novel bacteria, the microbiomes of different body sites retain distinct compositional profiles and display stability over time (Caporaso et al. 2011, Oh et al. 2016). Thus, most bacterial populations introduced into the human-associated microbiome also fail to establish.

Ecological studies have demonstrated that the success of invading taxa depends strongly on the biotic interactions within the resident community (Lodge 1993, Fey and Herren 2014). This is also observed in the context of the gut microbiome, where interactions take the form of exchanging and competing for metabolites (Kinnunen et al. 2016, Mullineaux-Sanders et al. 2018). For example, a recent study demonstrated that an alteration to the resident gut microbial community allowed invasive *Salmonella* to thrive on a newly abundant metabolite (Gillis et al. 2017). Thus, the resident gut community was indicative of resource availability, which mediated colonization resistance against *Salmonella*. Despite the importance of metabolite-mediated bacterial interactions, few models of microbial community invasion have explicitly included metabolites.

In this study, we investigate the role of the resident gut microbiome community in mediating resistance to invasion. First, we developed a computational model to study how metabolite cross-feeding and competition mediate success of an invading microbe. We then introduce a novel statistical method to identify invasive taxa in empirical microbial communities, and use this approach to study the invasive bacteria in three long-term gut time series. The gut microbiome is an ideal system to test ecological invasion theory, because the gut is at constant risk of invasion from consumed bacteria and from opportunistically invasive commensal bacteria (Benjamin et al. 2013). Finally, we evaluated whether gut connectivity could predict susceptibility to invasion by the pathogen *Clostridium difficile* in patients who received fecal microbiota transplant (FMT) therapy. We chose *C. difficile* as a test case because it is a well-studied bacterial pathogen whose colonization depends on the resident gut microbiome (Schubert et al. 2015).

## Methods

### Simulation Model

We constructed a mathematical model consisting of resident taxa, an invading taxon, and the metabolites required for cell reproduction. Taxa interact through competition for metabolites in the environment and through cross-feeding of metabolites. Of all possible metabolites in the model (*m*), each taxon required a randomly assigned unique subset of *n* metabolites for growth, giving each a distinct niche. Each taxon also excreted a non-overlapping subset of *q* metabolites. Excretion profiles were not necessarily unique. Cross-feeding, defined as direct flow of metabolites from one taxon to another, was possible if one taxon excreted a certain metabolite required by a different taxon. Other parameters in the simulation model included the proportion of possible metabolite exchanges due to cross-feeding that are realized (*p*), a competition coefficient (*c*), variability in competition coefficients among taxa (*v*), an input rate for metabolites (*i*), a flushing rate for metabolites and cells (*f*), metabolite input rates (*i*) and the number of taxa present in the community at the start (*x*). During each run, all possible combinations of *m* choose *n* metabolite requirement profiles were generated, and a random subset of *x* requirement and excretion profiles were assigned to the taxa. One of the *x* taxa had metabolic requirements matching the input metabolites. From the requirement and excretion profiles, every possible one-way metabolite flow was identified. A random subset of these possible exchanges were selected as realized cross-feeding relationships. Competition coefficients for each taxon were drawn from a normal distribution with mean *c* and a standard deviation of *v*.

In each time step, metabolites enter the environmental pool (Fig. 1a). Taxa then compete for these metabolites, with uptake rates governed by their competition coefficients, which quantify scavenging efficiency. Demand for each metabolite is calculated as the number of individuals lacking the metabolite multiplied by their respective competition coefficients. If total demand is greater than the available metabolites, metabolites are allocated among taxa in proportion to the demand of each taxon. We assume for simplicity that metabolite uptake amongst individuals in a population is arranged to maximize biomass production (Klitgord and Segrè 2011). Individuals that obtain one unit of each necessary metabolite reproduce and excrete their given metabolites. If these individuals were from taxa participating in cross-feeding, the excreted metabolites were preferentially available to the recipient taxon before entering the environmental pool. If the growing taxon had more than one exchange (i.e. more than one cross-feeder), an equal amount of metabolites were made available to each recipient taxon. Finally, a proportion *f* of individuals and environmental metabolites were flushed from the system.

**Figure 1:**
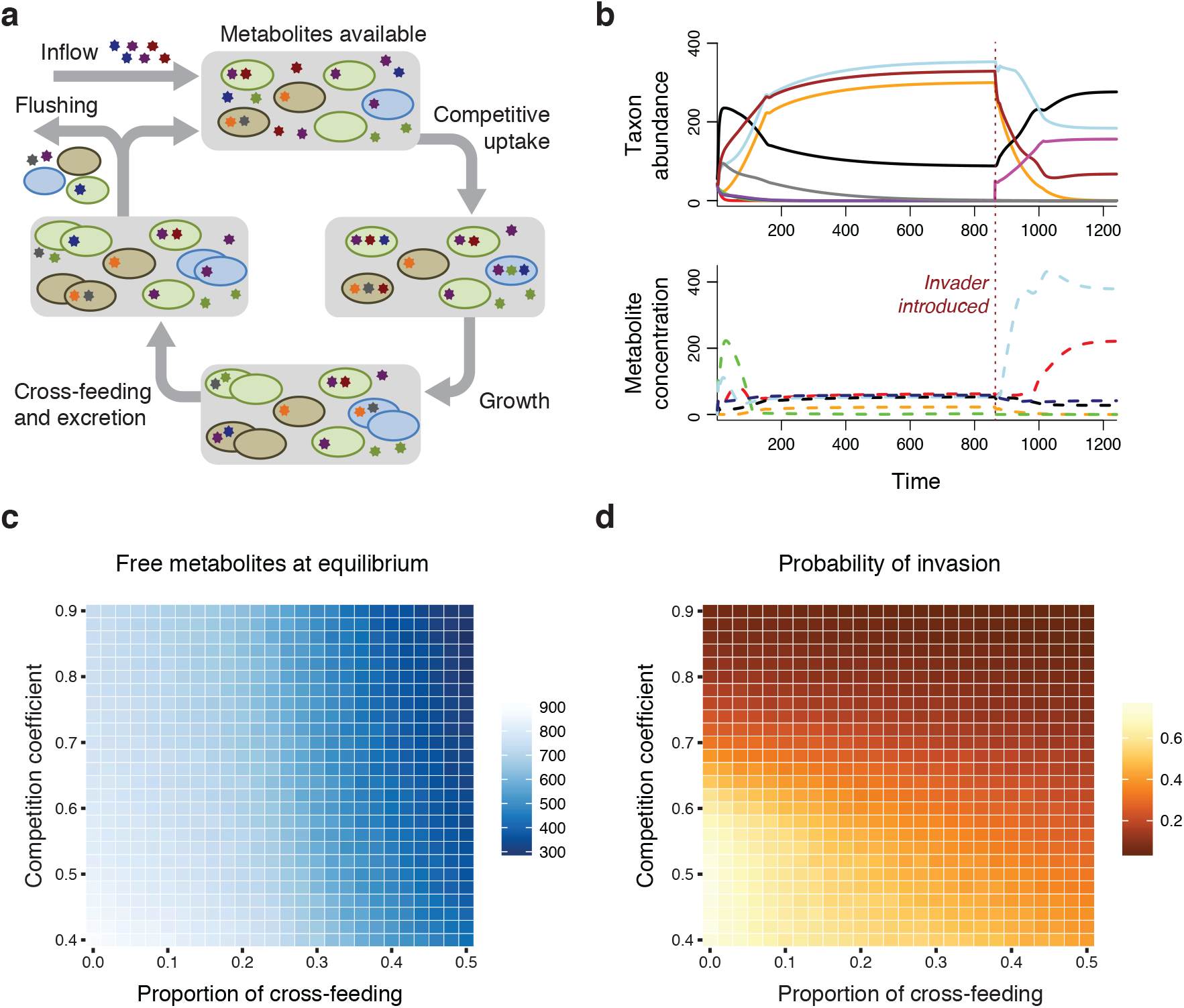
Stronger biotic interactions lead to more complete metabolite uptake and a decreased chance of invasion. a) A conceptual diagram showing the flux of metabolites during each step in the simulation model, beginning with metabolite inflow and ending with flushing. In this scenario, cross-feeding occurs from the blue cells to brown cells and from brown cells to both green cells and blue cells. b) The outputs of the simulation model are the taxon abundances over time and the environmental metabolite concentrations over time. The red dashed line shows the time point where the invader (pink) was introduced. In this case, the invader persisted in the community. c) Environmental metabolite concentrations at model equilibrium decrease as either parameter of biotic interaction (proportion of cross-feeding or competition coefficient) becomes stronger. The figure shows median values of 5000 runs for each parameter combination. d) The probability that an invader persisted in the community decreased when either parameter of biotic interaction (proportion of cross-feeding or competition coefficient) was stronger.

The invader was introduced after the community of resident taxa equilibrated (Fig 1b). The invader had a fixed competition value (in these simulations, 0.9), and participated in no cross-feeding. We reasoned that cross-feeding exchanges often need time to develop (e.g. time for proper spatial configuration [Pathak et al. 2012], construction of nanotubes [Pande et al. 2015], or within-host coevolution [Foster et al. 2017]), and that an invading taxon would therefore have no preexisting cross-feeding relationships. The run was completed once the model reached equilibrium after the invader was introduced. The outcomes recorded were persistence of the invader, standing pools of metabolites, total number of individuals in the community, and number of taxa persisting. We evaluated these outcomes while changing the strength of biotic interactions (magnitude of competition and proportion of cross-feeding).

For simulations analyzed here, we changed the mean strength of competition (*c*) between 0.4 and 0.9 and the proportion of cross-feeding (*p*) between 0 and 0.5. The standard deviation of competition values (*v*) was equal to 0.3 ^*^ *c*. Any randomly generated competition values below 0.01 were set to 0.01. We initialized the model with 15 taxa, 7 possible metabolites, 4 metabolites required for growth, 2 metabolites excreted, and an input rate of 100 units of each of 4 metabolites. The flushing rate was 0.15 during each time step. We used 5000 runs for each combination of competition coefficient and cross-feeding proportion. The model was determined to be at equilibrium when the maximum abundance change of any taxon was lower than 0.01. In a small fraction of model runs, the model resulted in a stable limit cycle (see SOM). In this case, the final values were recorded at 20,000 time steps.

### Dynamics of invasion in healthy subjects

Next, we present a technique to identify invasive taxa in three long-term time series of the human gut microbiome. Invasive taxa are, by definition, newly introduced into communities during discrete events. Thus, the distribution of invasive taxa over time shows clusters of presences and absences. We quantified the degree of presence/absence clustering using an equation that calculates the probability that a streak of successes would be observed in a series of Bernoulli trials (Feller 1968). This method of identifying invasive taxa avoids problems associated with defining invasive taxa as those newly observed in a community (Kinnunen et al. 2016). In that case, the taxa classified as invasive would change based on the reference time frame.

For each taxon, we identified the longest consecutive number of days in which the taxon was present (termed a “streak”). We also calculated the fraction of total samples in which was present. From these two values, it is possible to calculate a p value that described the *probability that a streak equal to or longer than the observed streak would occur if presences and absences were randomly distributed*. This probability (*qn*) is given by (see Feller 1968, p. 325 for a derivation):

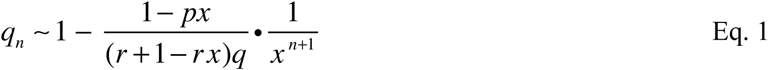

Where *n* is the number of samples, *r* is the length of the longest streak, *p* is the fraction of presences across the time series, *q* = 1 – *p*, and *x* is the root nearest to 1 (but not 1/*p*) of the equation:

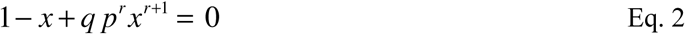

Using this formula, a small *q_n_* (p value) indicates that the observed streak of persistence is longer than would be expected; this denotes a highly patchy presence/absence distribution. We classified invasive taxa as those that were significantly patchy in their distribution (p < 0.01) and which were present in fewer than 50% of samples. We note that a cutoff of p < 0.05 may not be sufficiently conservative, due to expected autocorrelation in abundances. The presence threshold was necessary because common taxa that became temporarily absent during a disturbance would also have a significantly patchy distribution. Analyses were robust to changes in these cutoffs (see SOM). To account for differences in detection limits between samples due to differing sequencing depth, we set a universal detection limit across the entire dataset (see SOM). Any values lower than this limit were set to zero.

Data for human mirobiome subjects A and B were originally published in David et al. 2014. These data are near-daily stool samples from two unrelated male subjects. We downloaded the operational taxonomic unit (OTU) tables generated from closed reference OTU picking from the Qiita portal (ID 2196). We removed samples with fewer than 10000 reads and those with otherwise low detection ability (see SOM). We removed an additional two samples from subject B (days 252 and 318) because sampling became infrequent after day 242. This resulted in 321 samples from subject A and 182 samples from subject B. Data from subject C were obtained from the Qiita portal (ID 11052). Again, we used the OTU tables generated from closed reference OTU picking. We removed samples with fewer than 5000 reads and further removed OTUs with fewer than 10 reads across the full dataset. We subset the data to fecal samples from subject M03, and analyzed the longest window of consecutive fecal samples. This window corresponded to the 511 samples taken from 14 July 2015 to 06 February 2017. Four additional samples were removed due to poor detection limits.

### Calculation of connectivity using the cohesion statistics

We quantified connectivity of the resident gut community using cohesion statistics (Herren and McMahon 2017, https://github.com/cherren8/Cohesion), which give instantaneous measures of the degree of interconnectedness among OTUs in a microbial sample. Connectivity is hypothesized to arise from biotic interactions (see Herren and McMahon 2017 for further discussion). We operationally define connectivity as the true network of correlations between microbes, whereas cohesion is an estimate that quantifies connectivity.

In brief, cohesion statistics represent the average pairwise correlations between individuals in a sample, after correcting for methodological biases. There are two cohesion values for each sample, corresponding to connectivity from positive relationships and connectivity from negative relationships. The first step in the workflow is to calculate connectedness values for each taxon. Connectedness values are the average positive and negative correlation of the focal taxon with other taxa, after accounting for bias introduced by the compositional nature of the data. Cohesion values are weighted sums of the connectedness values multiplied by the abundances of the taxa in each sample. Taxa that are below specified abundance or persistence thresholds are not included in calculating cohesion, and are given connectedness values of zero.

For the long-term gut time series, we first-differenced the data before calculating cohesion values. This is a suggested step in the cohesion workflow to handle autocorrelation in abundances due to high sampling frequency (Herren and McMahon 2017). We set the persistence threshold for *inclusion* at 0.5 for the long-term gut data, which was the same cutoff for *exclusion* for defining invasive OTUs. Using the same cutoff for these two analyses meant that no taxon in the invasion analyses contributed to the cohesion values, thereby eliminating possible artificial relationships between the predictor and response variables in the multilevel model (see next section). We tested two possible mean abundance cutoffs of 0.001 and 0.0001. We evaluated which cutoff yielded a better fit of the population dynamics of invasive OTUs and used that cutoff value. For subjects A and B, we used a mean abundance cutoff of 0.001, while for subject C we used 0.0001.

### Multilevel model analysis and model selection

To evaluate whether connectivity could predict changes in taxon abundances, we built a hierarchical linear mixed effects model (hereafter called a “multilevel model”). We chose a multilevel model because it enables analysis of rare taxa without risk of over-parameterization. Invasive taxa are sufficiently rare that confident parameter estimates cannot be obtained for each taxon; instead, this type of analysis pools low-confidence estimates from each taxon to obtain a single overall estimate of how a predictor variable affects all invasive taxa. A template and sensitivity analysis of this type of analysis can be found in Jackson et al. 2012.

The predictors included in the multilevel models were fixed effects for the natural log-transformed abundance at time *t*, positive cohesion values of the communities at time *t*, and negative cohesion values of the communities at time *t*. The response variable was the natural log-transformed abundances of invasive taxa at time *t + 1*. We did not include time points when the OTU was absent (abundance = 0) at time *t* or at time *t +1*. This was for two reasons: 1) model residuals were improved by preventing zero-inflation, and 2) it was then possible to log transform the abundances without adding an arbitrary positive value. Random effects were included to allow mean abundance (intercept) to vary by OTU, for the effect of positive cohesion to vary by OTU, and for the effect of prior abundance to vary by OTU (effectively assuming density dependence varies by OTU). We did not include a random effect to allow negative cohesion to vary by OTU, because this random effect showed high collinearity with the random effect for positive cohesion, and negative cohesion was a weaker predictor than positive cohesion. We evaluated significance of random effects by comparing AIC values of models with and without the terms included.

In order to ensure that any significant results of our analyses of invasive OTUs were not spurious or statistical artifacts, we analyzed paired models of OTUs that were uncommon but non-invasive. We expected that OTUs classified as non-invasive would have a lesser response to cohesion values. Instead of analyzing OTUs with significantly patchy distributions (p < 0.01), we analyzed OTUs that showed no trend toward patchiness (p > 0.3) that were present in fewer than 50% of samples. We fit the same multilevel model for these OTUs. For subject A, highly collinear random effects caused convergence problems, so we removed the random effect that allowed density dependence to vary by OTU. A full table of results from the six multilevel models can be found in SOM.

### Relapse in patients treated with FMT

Finally, we tested whether the observed relationship between cohesion and resistance to invasion could be used to predict patient responses to a therapeutic microbiome intervention. We obtained data from a clinical study evaluating the role of the gut microbiome during FMT for the treatment of recurring *C. difficile* (Khanna et al. 2016). We downloaded the data from the Qiita portal (id 10057), again using the OTU table generated from closed reference OTU picking. As before, bacterial community composition data were obtained using 16S rRNA amplicon sequencing. We also contacted the original authors to ensure accurate interpretation of the metadata. FMT is a commonly used treatment for chronic *C. difficile* infection that has a higher success rate than antibiotic treatment (van Beurden et al. 2017). This study collected stool samples from 38 patients treated with FMT. Of these 38 patients, 29 had stool samples analyzed at day 28 post-FMT. The study also recorded whether patients experienced another *C. difficile* infection within two years after FMT. We hypothesized that patients who would later relapse would have lower community connectivity after FMT. To test this hypothesis, we calculated OTU connectedness values from the healthy donors, which we used to obtain cohesion values for the patients at 28 days post-FMT. The cutoffs for persistence and mean abundance of OTUs included in the cohesion calculation were 0.6 and 0.001. Because of the different variances of cohesion values among relapsing and cured patients, we used the non-parametric one-tailed Wilcoxon rank test for this analysis.

### Software used for analyses

All analyses and simulations were conducted in R, version 3.4.0. In the metabolite exchange simulations, taxon metabolite requirements were generated using the combinat package. The packages abind, data.table, and dplyr were used to manipulate matrices. Linear mixed effects models were fit using the lme4 package. Conditional R^2^ values were obtained using the MuMIn package. Significance values for models were obtained using the lmerTest package.

## Results and Discussion

### Simulation model

Simulation results showed that the probability that an invader could persist in the community was strongly related to the strength of biotic interactions in the resident community. As the strength of either interaction (competition or cross-feeding) increased, standing metabolite pools decreased (Fig. 1c). Similarly, invaders were less successful when interactions were stronger (Fig. 1d). The two types of interactions had different mechanisms of reducing metabolites available to the invader. Stronger competition resulted in greater metabolite scavenging from the environment by resident taxa, thereby decreasing the probability that the invader could acquire metabolites. Conversely, cross-feeding decreased the amount of excreted metabolites entering the environmental pool due to direct provisioning of metabolites. Thus, stronger biotic interactions, whether originating from competition or cross-feeding, also led to lower rates of invasion. Our model supports the hypothesis that bacterial interactions can mediate invasion by inducing resource scarcity, thereby making it difficult for invaders to gain a foothold (Kinnunen et al. 2016).

In the absence of cross-feeding, the model simplifies to the well-known result that only *m* taxa can persist on *m* metabolites (Tilman 1982). When cross-feeding was introduced, one taxon for every niche ( *m* choose *n* taxa) could be present. Interestingly, this strong increase in diversity when allowing for cross-feeding is another possible explanation for the paradox of the plankton (Hutchinson 1961). The model has a carrying capacity determined by *i*, *n*, *q*, and *f*. If metabolite exchange from cross-feeding were configured such that every excreted metabolite were immediately consumed, the maximum community density (carrying capacity) is equal to

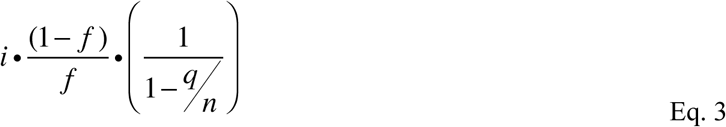

### Dynamics of invasion in healthy subjects

Next, we used our new approach for identifying invasive taxa in three long-term time series of the human gut microbiome. We designated taxa with significantly temporally clustered presence-absence profiles (p < 0.01) as invasive (Fig. 2).

**Figure 2:**
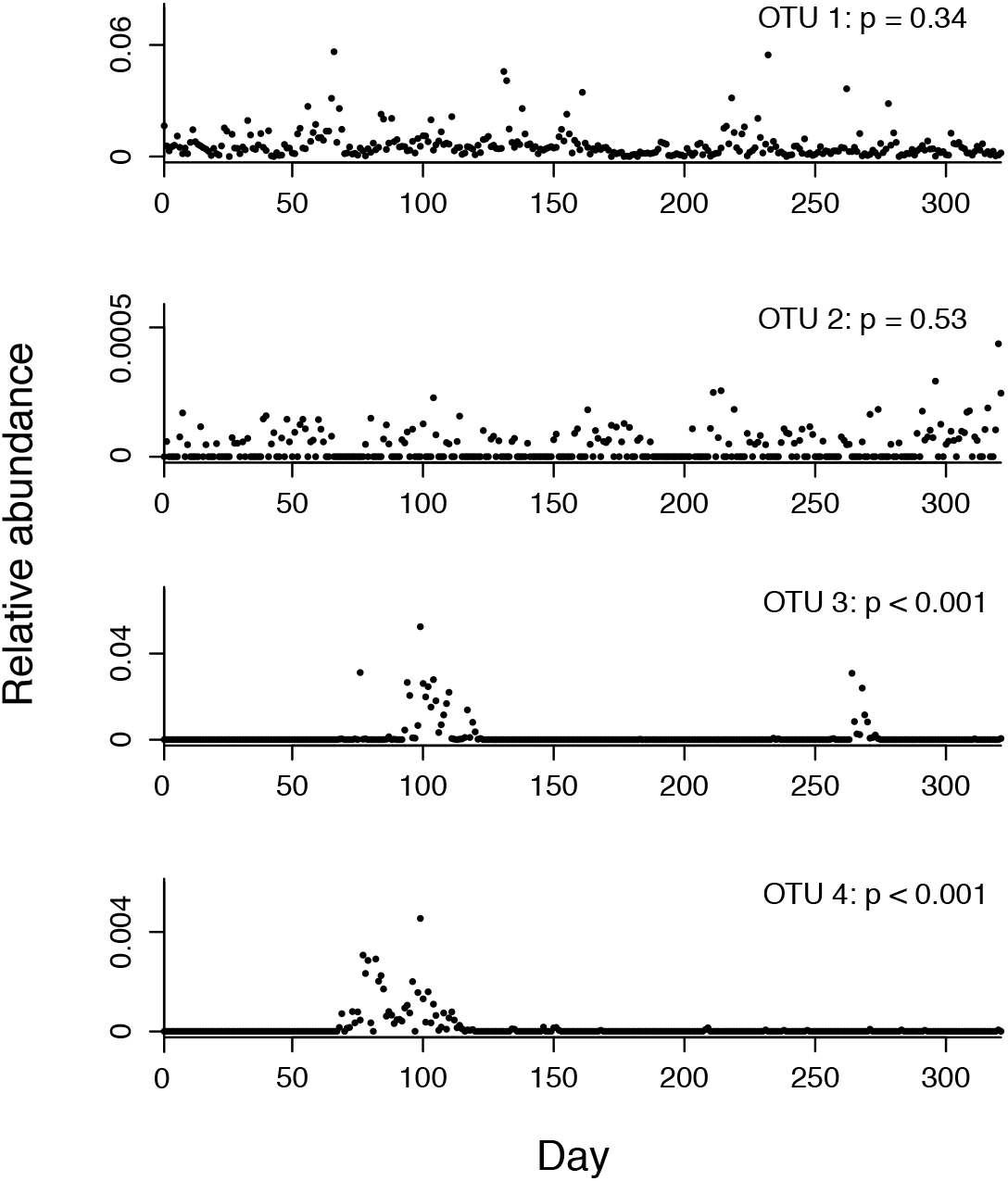
Invasive taxa have distinct time-series signatures. We classified each OTU as invasive or non-invasive based on its presence/absence distribution across the time series. We used a formula for quantifying the temporal clustering of presence/absence data to make this classification (Feller 1968). This formula calculates the probability of observing consecutive presences of a taxon, which we call a “streak”. The p values displayed correspond to the probability that the OTU’s longest streak would occur if presences and absences were distributed independently across the time series. Taxa showing interspersed presences and absences across the time series were categorized as not significantly clustered, and therefore non-invasive (OTU 1 and OTU 2). Taxa showing highly segregated presences and absences across the time series were categorized as significantly clustered, and therefore invasive (OTU 3 and OTU 4).

Invasive taxa were common in all three subjects. The total number of OTUs in each dataset was 1828 for subject A, 1771 in subject B, and 4918 in subject C. In subject A, we identified 250 invasive OTUs (subject B: 441, subject C: 131) that comprised an average of 2.2% (subject B: 8.6%, subject C: 2.9%) of the gut bacterial community. Although invasions were common, invader abundances were low and the duration of invasions were short. Median peak invader abundance was 0.053% (subject B: 0.047%, subject C: 0.13%) of the community, with an interquartile range of 0.022–0.24% (subject B: 0.022–0.15%, subject C: 0.058 – 0.32%), and median invasion duration was 7 days with an interquartile range of 4-14 days (subject B: median 8 days, IQR of 4–15 days, subject C: median 7 days and IQR of 5–14 days). The large number of invaders in subject B is likely explained by this subject’s gastrointestinal infection during the course of the sampling period.

The multilevel models showed that cohesion values were significant predictors of abundance changes for invasive OTUs, but not for non-invasive OTUs. In all subjects, invasive OTUs declined in abundance when positive cohesion in the gut community was stronger (subject A: p < 10^−8^, slope 95% confidence interval = [−8.32, −4.24]; subject B: p < 10^−10^, CI = [−8.32, −4.59]; subject C: p < 10^−5^, CI = [−12.3, −4.98]; Fig. 2). Please see SOM for full results tables. Invasive OTUs in subjects A and B showed no response to negative cohesion, while invasive OTUs in subject C declined as negative cohesion became stronger (subject A: p = 0.18, CI = [−2.27, 0.45]; subject B: p = 0.10, CI = [−1.72, 0.16]; subject C: p < 10^−4^, CI = [2.28, 6.74]). Conversely, OTUs that were not classified as invasive showed no response to changes in cohesion in subjects A (positive: p = 0.11, CI = [−6.07, 0.70]; negative: p = 0.85, CI = [−2.56, 3.08]) and B (positive: p = 0.12, CI = [−8.92, 1.09]; negative: p = 0.57, CI = [−2.66, 4.81]), and showed increases in abundance in subject C (positive: p = 0.014, CI = [2.56, 24.7]; negative: p = 0.068, CI = [−13.7, 0.63]). The range of positive cohesion values observed in communities from subject A was 0.035 to 0.148 (subject B: 0.026 to 0.121, subject C: 0.024 – 0.114). A decrease in cohesion of 0.05 was associated with an increase of the average invader’s abundance by 37% (subject B: 38%, subject C: 54%) (Fig. 3). The conditional R^2^ values for the hierarchical models of invasive taxa were 0.45 for subject A, 0.61 for subject B, and 0.44 for subject C.

**Figure 3:**
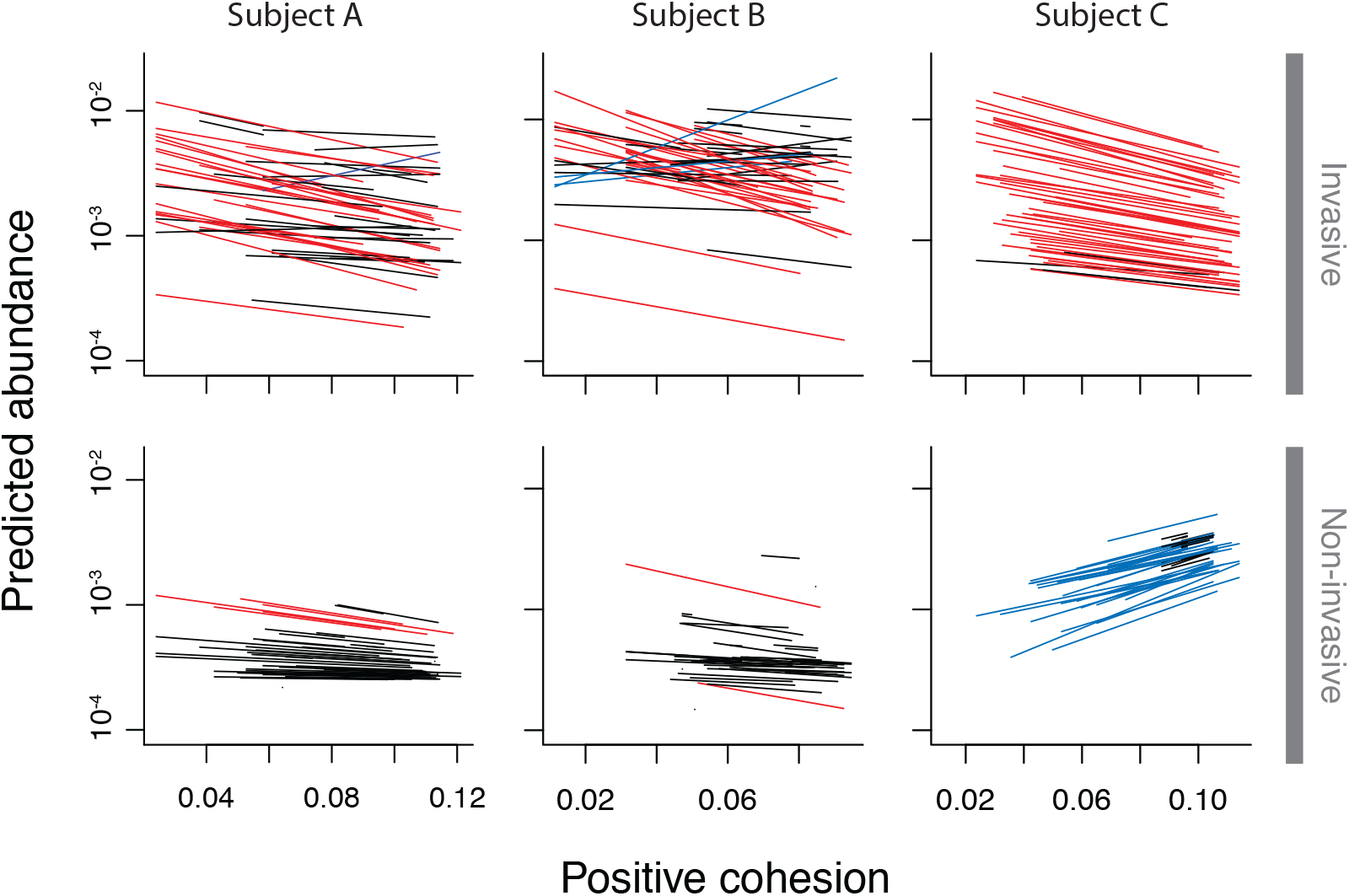
Invasive OTUs decrease in response to stronger cohesion, whereas non-invasive OTUs do not. Each panel shows the predicted abundances of the 50 most abundant taxa in each multilevel model analyzing OTU abundance as a function of cohesion. Each line indicates the predicted abundance of the OTU at time *t + 1* if the OTU were observed at a relative abundance of 0.01 at time *t*. Red lines indicate taxa that decreased by at least 50% over the range of observed positive cohesion values. Blue lines indicate taxa that increased by at least 50%. Although there was variability in how the OTUs responded to positive cohesion, invasive OTUs had a significant overall negative response to stronger positive cohesion in all three subjects (top row). For comparison, we also analyzed OTUs that were uncommon but non-invasive. In subjects A and B, these OTUs showed no significant response to cohesion, whereas OTUs in subject C the non-invaders increased in response to positive cohesion (bottom row).

### Relapse in patients treated with FMT

Finally, we analyzed the data from a clinical study evaluating the role of the gut microbiome during FMT for the treatment of recurring *C. difficile* (Khanna et al. 2016). Stool samples were taken from patients before and after FMT, as well as from donors. We found that patients who would have a future relapse within 2 years after FMT had weaker gut cohesion at 28 days post-FMT than patients who were cured (Fig. 4). Signed Wilcoxon rank tests were significant for both positive cohesion (p = 0.012) and negative cohesion (p = 0.015). Thus, weak connectivity of the patient gut bacterial community after FMT was an indicator that the patient was more likely to relapse. Prior efforts to predict *C. difficile* relapse from the microbiome did not find post-FMT indicators of relapse, although connectivity was not previously considered (Pakpour et al. 2017).

**Figure 4:**
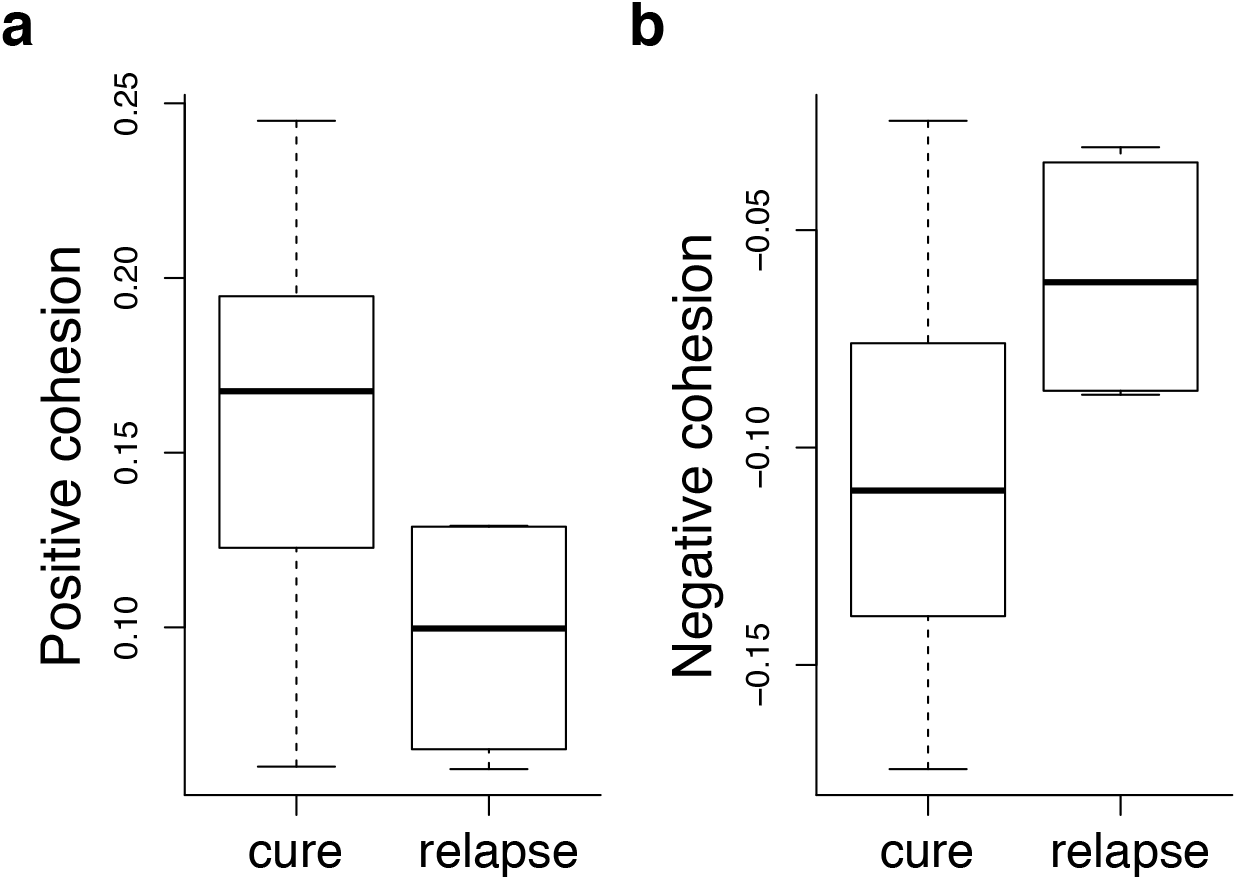
Resident microbiome cohesion predicts *C. difficile* relapse in patients treated with FMT. Patients who would later experience a relapse of *C. difficile* had lower gut bacterial cohesion values at 28 days post-FMT. We calculated connectedness values of OTUs in healthy stool samples, which were then used to calculate cohesion values for patients after they received FMT. Relapsing patients had weaker positive cohesion (Wilcoxon signed rank test p value = 0.012) and weaker negative cohesion (Wilcoxon signed rank test p value = 0.015) after treatment.

We conducted similar analyses using other possible predictors of relapse to evaluate the comparative explanatory power of the cohesion statistics. Other potential predictors of relapse were not statistically significant, including post FMT gut diversity (p = 0.92 using a one-way ANOVA), Bray-Curtis dissimilarity from the donor community (p = 0.69 using a one-way ANOVA), and cohesion of the donor community (p = 0.18 for positive cohesion, p = 0.14 for negative cohesion using signed Wilcoxon rank tests). The small number of relapsing patients is expected, because the success rate of FMT is high. However, when using cohesion values as the predictor, the effect size was sufficiently large that the small sample size did not impede identifying significant separation between the cured and relapsing groups.

### Incorporating invasion into ecological models of the gut microbiome

The three cases considered here consistently showed that stronger connectivity of the resident bacterial community led to increased colonization resistance. Agreeing with ecological invasion theory, most bacterial invasions were short and ultimately unsuccessful. Prior studies have hypothesized that antibiotics increase susceptibility to invaders by eliminating resident microbes (David et al. 2014, Schubert et al. 2015). We further suggest that the most highly connected taxa are disproportionately important to maintaining resistance against invaders, and that successful gut microbiome interventions (such as probiotics [Johnston et al. 2016] or dietary changes [Griffin et al. 2017]) may be attributable in part to restoring connectivity among resident gut microbes.

Interestingly, our results show predictive power independent of the role of the host immune system, which is a substantial contributor to host susceptibility to invasion (Round and Mazmanian 2009). Observing the responses of each invasive gut OTU (Fig. 3), it is clear that some invasive taxa had a positive or neutral response to increased gut connectivity. The variability of invader response to cohesion may be explained by factors not considered in this analysis. For example, it is possible that the taxa that are largely unaffected by cohesion are instead regulated by an immune response or the synthesis of antimicrobial compounds (Mullineaux-Sanders et al. 2018). Thus, we expect that resident community connectivity is only one of several factors affecting gut invasion. Furthermore, although the empirical results are consistent with the interpretation that pairwise correlations reflect biotic interactions, we cannot exclude the possibility that other factors, such as environmental forcing, contributed to cohesion values.

The mechanisms by which FMT resolves *C. difficile* infection are still poorly known (Bajaj et al. 2017, Patron et al. 2017, Zuo et al. 2017). Indeed, when announcing the FMT National Registry, the American Gastronomical Society wrote that the widespread use of FMT “has advanced the practice of gut microbiota manipulation in patients more rapidly than our scientific understanding” (Kelly et al. 2017). Our results indicated that FMT is more successful when post-transplant communities have many highly connected taxa. We therefore hypothesize that successful intervention via FMT reestablishes microbial interactions, thereby creating a microbial community that can repel invaders. Taken together, our analyses show that strong connectivity of the resident gut bacteria appears to be an important indicator of a resilient microbial community.

